# Complement-dependent synapse loss and microgliosis in a mouse model of multiple sclerosis

**DOI:** 10.1101/720649

**Authors:** Jennetta W. Hammond, Matthew J. Bellizzi, Caroline Ware, Wen Q. Qiu, Priyanka Saminathan, Herman Li, Shaopeiwen Luo, Yuanhao Li, Harris A. Gelbard

## Abstract

Multiple sclerosis (MS) is an inflammatory, neurodegenerative disease of the CNS characterized by both grey and white matter injury. Microglial activation and a reduction in synaptic density are key features of grey matter pathology that can be modeled with experimental autoimmune encephalomyelitis (EAE). Complement deposition combined with microglial engulfment has been shown during normal development and in disease as a mechanism for pruning synapses. We tested whether there is excess complement production in the EAE hippocampus and whether complement-dependent synapse loss is a source of degeneration in EAE using C1qa and C3 knockout mice. We found that C1q and C3 protein levels were elevated in EAE mice. Genetic loss of C1qa provided a small degree of protection from EAE-induced decreases in synaptic density. However, C1qa knockout EAE mice had similar levels of microglial activation and identical clinical scores as WT EAE mice. C3 knockout mice were largely protected from EAE-induced synapse loss and microglial activation, results that correlated with a reduction in the EAE clinical score. Thus, pathologic expression and activation of the early complement pathway drives a portion of the synapse elimination observed in the EAE grey matter.

## Introduction

Multiple sclerosis is an immune-mediated disease of the CNS involving damage to both the white and grey matter. Current immunomodulatory therapies for MS are effective at minimizing relapses and white matter lesion burden, but are less effective at preventing progressive neurodegeneration and cognitive deficits [1]. Grey matter degeneration is highly correlated with the progressive component of MS and its accompanying physical and cognitive disabilities [2–5]. Grey matter pathology involves focal demyelinated lesions and widespread atrophy of cortical and subcortical regions. MS demyelinated grey matter is characterized by activated microglia, axon damage, neuron and glial cell loss, and decreased synaptic density [6–8]. Interestingly, activated microglia and synapse loss also occur in myelinated areas of normal-appearing grey matter [9, 10] suggesting direct synaptic injury associated with activated microglia may be involved.

Microglia have been shown to prune synapses by phagocytosis during development by a process of complement-mediated opsonization and subsequent phagocytosis [11–14]. The complement system can be activated by deposition of C1q, which is thought to tag synapses for elimination and triggers the classical complement pathway, or via the alternative and lectin pathways that are independent of C1q. All three initiating pathways converge to cleave C3 to C3a, a soluble chemokine that recruits phagocytic cells, and C3b which is deposited on activating surfaces as an opsonin that facilitates phagocytosis via the microglia-specific complement receptor 3 (CR3) [11, 15]. C3b can also initiate the lytic pathway leading to formation of the membrane attack complex (MAC) that causes cell lysis, but it is the early complement pathway (C1q - C3) that is implicated in synapse pruning by microglia [11, 16, 17].

While critical for development, complement is also recruited to mediate pathologic synapse loss by microglia during aging as well as in diseases including frontotemporal dementia, Alzheimer’s disease, viral encephalitis, traumatic brain injury, and schizophrenia [16–22]. Yet, microglial activation and synapse loss may occur independent of complement in other diseases such as the mutant SOD1 model of ALS and a model of HIV associated neurocognitive disorder (HAND) [23, 24]. Thus, the goal of this study was to evaluate whether complement-dependent synapse loss contributes to grey matter degeneration in EAE, which may also provide insight into its role in MS..

Complement involvement in the white matter pathology of MS is well-established. Complement activation products including the MAC complex are abundant in human white matter lesions [25, 26]. Evaluation of the role of the terminal lytic pathway in EAE has yielded mixed results [27–32]. However, the alternative complement pathway plays a major role in driving pathology in EAE. Knockout of C3, Factor B, or receptors to C3 cleavage products (C3aR, CR3, and CR4) all reduce white matter lesion pathology and motor impairment measured by the EAE clinical score [33–38].

The role of complement in MS grey matter degeneration has only begun to be addressed. Two recent studies on human postmortem samples indicate that early complement components C1q, C3b-iC3b, C3d (a cleavage product of C3 that marks areas of chronic complement activity), and alternative pathway component Bb do accumulate in cortical, hippocampal, and thalamic grey matter of MS patients [9, 39]. Within the hippocampus, C1q and C3d were increased in both myelinated and demyelinated regions of CA1-3 that also displayed synapse loss and microglial activation. C1q and C3d also co-localized with synaptic markers within microglial processes, suggesting that complement-opsonized synapses were potentially phagocytized by microglia [9]. There was no sign of the terminal MAC in the hippocampus [9, 39], showing that the early complement pathway is the most relevant to progressive MS grey matter pathology. As the early complement pathways are correlated with the synapse reduction observed in MS patients, it is important to determine whether the relationship is causative and thus potentially preventable using animal models.

MOG_35–55_-induced EAE provides a good animal model of several aspects of MS grey matter injury. The hippocampi of EAE mice have pronounced microglial activation, synapse loss, and some atrophy of pyramidal cell layers that occurs independently of extensive myelin loss, axon injury, or significant accumulation of peripheral immune cells [40–45]. Chronic EAE mice, 66+ days post immunization, exhibit significant reductions in brain volume including atrophy of the hippocampus [46]. EAE mice also show impaired synaptic transmission and deficits in hippocampal-dependent behavior [41, 47–49] indicating that EAE mice have cognitive impairments analogous to progressive MS patients.

We have used the MOG_35–55_ EAE model to test the hypothesis that the EAE inflammatory environment results in excess complement production and activation in the brain to make synapses vulnerable to pruning from phagocytic microglia. Our results show that complement proteins C1q and C3/C3d are elevated in the hippocampus of EAE mice and that genetic deletion of C1q or C3 provides varying levels of protection from EAE-induced synapse loss and microglial activation.

## Methods

### Animals

Male and female C57BL/6 mice were obtained from Jackson Laboratories (Bar Harbor, ME) at age 7 weeks and were housed for at least 1 week prior to EAE induction. A C3 knockout breeder pair (with C57BL/6 background) was obtained from Jackson (Stock # 029661; [50]). A C1qa knockout (KO) breeder pair was obtained from Dr. Andrea Tenner (University of California, Irvine; [51]). C1q is an oligomeric protein complex originating from 3 genes: C1qa, C1qb, and C1qc. Loss of C1qa prevents correct folding and secretion of the C1q multimer with a complete loss of function [51]. Groupings of male and female mice were used for all reported experiments in relatively equal numbers. Two-way ANOVA tests showed no significant main effects of sex or significant interactions of sex with immunization status or genotype so the data was combined and reported from both sexes.

### EAE

We immunized mice with a pre-mixed emulsion (Hooke Labs EK-2110) containing myelin oligodendrocyte glycoprotein peptide, amino acids 35-55 (MOG_35-55_) in complete Freund’s adjuvant (CFA) containing heat-inactivated mycobacterium tuberculosis H37RA. A 100µl emulsion was injected subcutaneously at each of two sites over the upper and lower back. Sham-immunized sex-matched littermate controls were injected with control emulsion without MOG_35-55_ (Hooke Labs CK-21100), and otherwise received identical treatment. Mice of each genotype were randomly assigned to either the sham or EAE treatment groups. Mice were injected with pertussis toxin (Hooke Labs, 90-400ng intraperitoneally (i.p.), with the dose adjusted according to potency of each lot per manufacturer recommendations for each sex) on 0 and 1 day post-immunization (dpi). In one cohort, WT mice received ~100ul i.p. twice daily of Vehicle solution: 5% DMSO, 40% polyethylene glycol 400, and 55% normal saline beginning on the first day of visible motor deficits (EAE score 0.5 or greater) with timing matched for sham controls. All mice were monitored daily for signs of clinical disease and scored for motor deficits as follows: 0, no deficit; 0.5, partial tail paralysis; 1, complete tail paralysis; 2 hind-limb weakness; 2.5, paralysis of one hind limb; 3, paralysis of both hindlimbs; 3.5, hindlimb paralysis and forelimb weakness; 4; quadriplegia; 5; moribund.

### Western blots

At twenty-eight days post-immunization mice were perfused briefly with PBS, hippocampi were isolated, immediately frozen on dry ice, and stored at −80°C. Hippocampal biopsies were homogenized in RIPA lysis buffer containing a protease inhibitor cocktail (MilliporeSigma: Calbiochem, SetV). The cell lysate was kept on ice with periodic vortexing for 30 minutes then centrifuged at 13,000 rpm for 10 minutes and repeated with the resulting supernatant. The protein concentration of the final supernatant lysate was assayed with a detergent-compatible Bradford assay (Pierce). Equivalent amounts of protein were then mixed with loading dye and run on an 8 or 12% SDS-PAGE gel and transferred to PVDF. Membranes were blocked with 5% milk in TBSt for 30 minutes and probed with primary antibodies: C1q (kind gift of Andrea Tenner, 1151, [52]), C3d (R&D systems, AF2655), CFH (R&D systems, AF4999), and actin (Santa Cruz, SC-47778) in 5% milk in TBSt at 4 degrees overnight. Membranes were washed 3 times in TBSt and then incubated with HRP secondary antibodies for 1 hour in 5% milk in TBSt. After washing we applied ECL substrate (Pierce) and developed membranes using a digital imager (Azure Biosystems). Membranes were stripped using buffer consisting of 200mM glycine; 0.1% SDS, 1% Tween 20, pH 2.2 and re-probed up to 3 times. Western blot band densities were quantitated using Image J. Band densities for each animal were normalized to the control sham group mean from the same gel and graphed as mean ± SEM.

### RNA isolation and qPCR

At 28-30 days post-immunization mice were perfused briefly with PBS and hippocampi were isolated and flash frozen using dry ice. RNA from a single hippocampus per mouse was obtained using the Nucleospin RNA kit (Macherey Nagel; Cat# 740955). cDNA was synthesized using random hexamer primers and the Superscript III first-strand synthesis system for RT-PCR (ThermoFisher, Cat# 18080-051). qPCR was performed using Taqman probes: C1qa Mm00432142_m1; C3 Mm_00437838_m1; IPO8 Mm_01255158_m1 (ThermoFisher). Delta Ct values for C1q and C3 were calculated relative to the housekeeping gene IP08 for each sample and fold changes were calculated relative to sham controls.

To obtain RNA specifically from CD11b+ microglia/myeloid cells, mice were perfused with PBS 28-30 days post-immunization and the cortex and hippocampi were dissected and combined. Single-cell suspensions free from myelin and RBCs were obtained using the Adult Brain Dissociation kit (Cat# 130-107-677) and gentleMACS Octo Dissociator from Miltenyi according to the kit instructions. Single cell suspensions were incubated with CD11b magnetic beads (Miltenyi, cat# 130-093-636) then CD11b+ cells were separated from other cell types using a magnetic MACs separator and LS columns (Miltenyi). CD11b+ cells were pelleted and RNA isolation, cDNA synthesis, and qPCR reactions were processed similar to the whole hippocampal tissue outlined above.

### IHC

At 28-30 days post-immunization, mice were anesthetized with ketamine/xylazine (100 and 10 mg/kg, respectively) and intracardially perfused for 1 minute with phosphate-buffered saline (PBS) containing EDTA (1.5 mg/ml) followed by 4% paraformaldehyde (PFA) in PBS. Brains were post-fixed for 18-24 hours in 4% PFA then stored in PBS at 4°C. Brains were cut into 40μm-thick coronal sections using a vibratome (Leica V1000) and stored in a cryoprotectant mixture of 30% PEG300, 30% glycerol, 20% 0.1 M phosphate buffer, and 20% ddH_2_O at −20°C. We performed free-floating-section IHC. The sections were washed three times for 30 min in PBS to remove the cryoprotectant. Then sections were incubated in 100mM glycine in PBS for 30 minutes followed by citrate antigen unmasking at 37°C for 30 minutes (Vector, H3300 with 0.05% tween 20). Primary antibodies were diluted in blocking buffer consisting of 1.5% BSA (MilliporeSigma; A3294), 3% normal goat serum (Vector Laboratories; NGS; S1000) or 3% normal donkey serum (MilliporeSigma, 566460), 0.5% Triton X-100 (Promega, H5142), and 1.8% NaCl in 1× PBS. We used the following antibodies in these experiments: rabbit anti-Iba1 (Wako Biochemicals, 019-19741; 1:1000) rabbit anti-C1q (Abcam Ab182451 clone 4.8; 1:500), goat anti-C3d (R&D Systems, AF2655, 1:500), chicken anti-Homer1 (Synaptic Systems, 160006, 1:500), and mouse anti-PSD95 (NeuroMab, 75-028, 1:500). Sections were incubated in the primary antibody mixture for 2-3 days at room temperature with agitation. Sections were then washed three times for 30 min in 1× PBS with 1.8% NaCl and then incubated overnight at room temperature in Alexa Fluor-conjugated secondary antibodies (1:500; Invitrogen or 1:250 Jackson Immuno Research) in blocking buffer. Finally, sections were washed 3 times with 1×PBS with 1.8% NaCl, mounted on slides with Prolong Gold or Diamond mounting agents (Life Technologies; P36935 and P36961).

### Imaging

IHC sections were imaged with a Hamamatsu ORCA-ER camera on an Olympus BX-51 upright microscope with Quioptic Optigrid optical sectioning hardware with identical light intensity and exposure settings for all animals within each image set. The acquisition was controlled with Volocity 3DM software (PerkinElmer Life and Analytical Sciences). Objectives: 4x, 0.13 NA; 10x, NA 0.4; 60x NA 1.4. Image analyses was done using Volocity. Differences in *z*-axis registration of different fluors were corrected by calibration with multicolor fluorescent beads. For 60x images, we imaged the CA1-stratum radiatum hippocampal area collecting 6 image stacks (10 or 12μm thick; *z*-step = 0.3μm) usually from the left and right hemispheres of 3 separate sections. All image stacks are displayed as maximum intensity protections. IBA1+ cell morphology measurements (volume, surface area and skeletal length) were obtained using an intensity-based automated segmentation algorithm in Volocity (“find objects”) followed by a fine filter. Homer1 and PSD95 puncta were identified using the “find spots” algorithm in Volocity software, which identifies puncta based on local intensity maxima that also exceed an intensity threshold value. For all measurements, values from image stacks in 6 tissue sections per mouse were averaged and expressed relative to sham-immunized controls from the same genotype and graphed as mean ± SEM. Experimenters were blinded to genotype or immunization group throughout each IHC experiment and subsequent data analysis.

### Statistics

GraphPad Prism software was used to perform all statistics. *N* values (number of animals) for each experiment are reported in the figure legends and come from two or more independent cohorts. We defined significance as *p* <0.05. Statistical tests are listed in text and figure legends. In general Student’s t-tests were used for statistical analysis comparing sham vs EAE in WT-only data sets. ANOVAs followed by Sidak’s multiple comparisons test were used when comparing sham vs EAE affects in multiple genotypes. The non-parametric Kruskal-Wallis test with Dunn’s multiple comparisons was used for analysis of the clinical scores. All data are expressed as the mean ± standard error of the mean (SEM).

### Study approval

Animal care and use were carried out in compliance with the US National Research Council’s Guide for the Care and Use of Laboratory Animals and the US Public Health Service’s Policy on Humane Care and Use of Laboratory Animals. Protocols were approved by the University Committee on Animal Resources at the University of Rochester.

## RESULTS

### C1q and C3 expression in EAE hippocampus

Our lab has previously shown that EAE immunization results in significant synapse loss in the CA1-statum radiatum of the hippocampus, which is largely spared from demyelination [40, 53]. To determine if the EAE hippocampus also exhibits increased complement production and deposition in EAE that could make synapses vulnerable to phagocytosis by glia, we first analyzed C1q and C3 protein and mRNA expression by Western blot and qPCR. By western blot, found that C1q protein expression in the hippocampus of EAE mice was increased 2.6-fold compared to sham (Figure 1 A-B; p<0.001; *t*-test). Expression of full length C3 protein was also elevated in EAE mice 1.9-fold above sham controls (Figure 1 A-B; p=0.003; *t* test). qPCR analysis of mRNA expression from hippocampal tissue shows there is a significant increase in the local expression of both C1qa and C3 in EAE mice compared to sham controls (Figure 1 C; C1q: 2.1 fold above sham controls p<0.001 *t*-test; C3: 8.4 fold above sham controls p=0.001). Thus the increased protein expression of C1q and C3 in the EAE hippocampus is due at least in part to local gene expression and not simply due to blood brain barrier breakdown.

**Figure 1:**
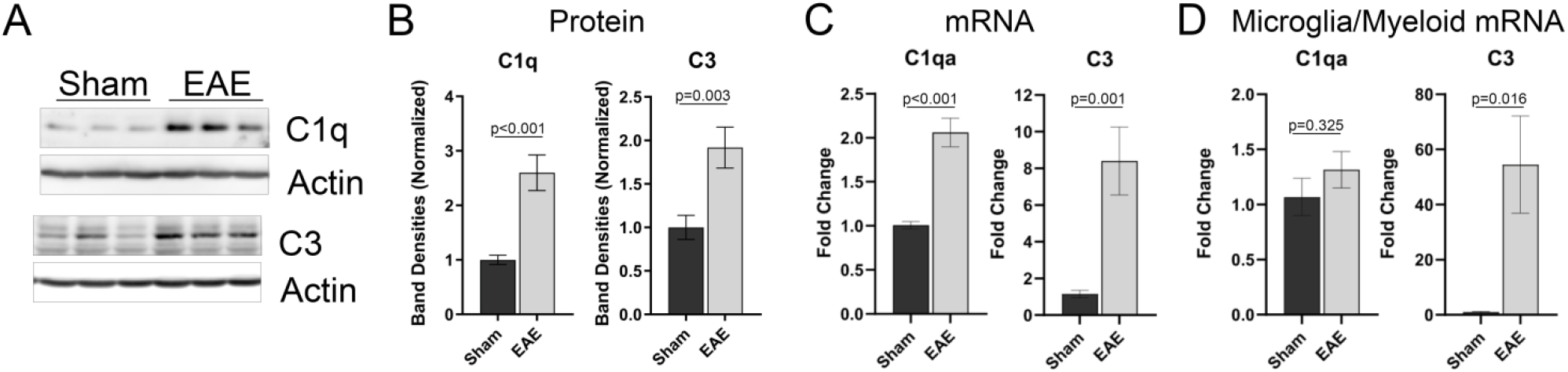
EAE mice have increased expression of complement components C1q and C3 in the hippocampus. Hippocampi were isolated from Sham or EAE mice at 25-30 days post immunization. A-B) Hippocampal lysates were probed by western blot for expression of C1q, C3, and Actin. A) Gel images show lysates from 3 representative animals per group. B) Quantification of western blot band densities for C1q and C3 have n=7-11 animals per group. All band densities are normalized to sham controls. C) RNA isolated from the hippocampus of sham or EAE mice was transcribed into cDNA and analyzed by qPCR for expression of C1qa and C3 with fold changes relative to sham controls. n=10 per group. D) qPCR results of C1qa and C3 gene expression from CD11b+ microglia/myeloid cells isolated from the hippocampus and cortex of sham or EAE mice. n=5 per group. For all graphs: Error bars=SEM. Statistical Analysis: Student’s t-test.

Because microglia are thought to be the main producers of C1q in the healthy hippocampus [54] and can also upregulate C3 when activated, we used qPCR to analyze the local gene expression of the CD11b+ microglia/myeloid cells in sham and EAE mice. In order to obtain sufficient cells, we isolated CD11b+ cells from the hippocampus and cortex. We found that CD11b+ cells in EAE mice significantly overexpress C3 at 28-30 days post immunization compared to sham controls (Figure 1 D; 54.5 fold increase; p=0.016, *t*-test). However, there was only a small trend for increased C1qa expression by CD11b+ cells in EAE but no significant difference (Figure 1 D; 1.3 fold p=0.32 *t*-test). This shows that microglia (and other myeloid cells) contribute to the elevated C3 expression and perhaps some C1q expression in EAE. However, we cannot rule out EAE induced elevated expression of complement proteins by other cell types in the hippocampus.

Next, we performed IHC using anti-C1q and anti-C3d antibodies to see where the elevated C1q and C3 were located in the EAE hippocampus (Figure 2). By IHC we measured a 1.6-fold increase in mean C1q fluorescence across the hippocampus in EAE vs sham mice (Figure 2 A-B; p=0.02; *t*-test) with significant increases ranging from 1.5-1.9-fold in all sub-regions of the hippocampus (Figure 2 A, C). At high (60-100x) magnification we observed that C1q was diffusely localized throughout the neuropil but also was localized at higher density in small punctate regions, some of which co-localized with synapses or along the dendrites in both sham and EAE brains (identified by the postsynaptic marker PSD95) (Figure 2 D-E). However, we also observed abundant heterogeneous expression of C1q that did not co-localize to synapses, but may contact other structures in the neuropil (Figure 2 E). We additionally observed higher C1q expression around blood vessels (Figure 2 G right panels) in both sham and EAE mice.

**Figure 2:**
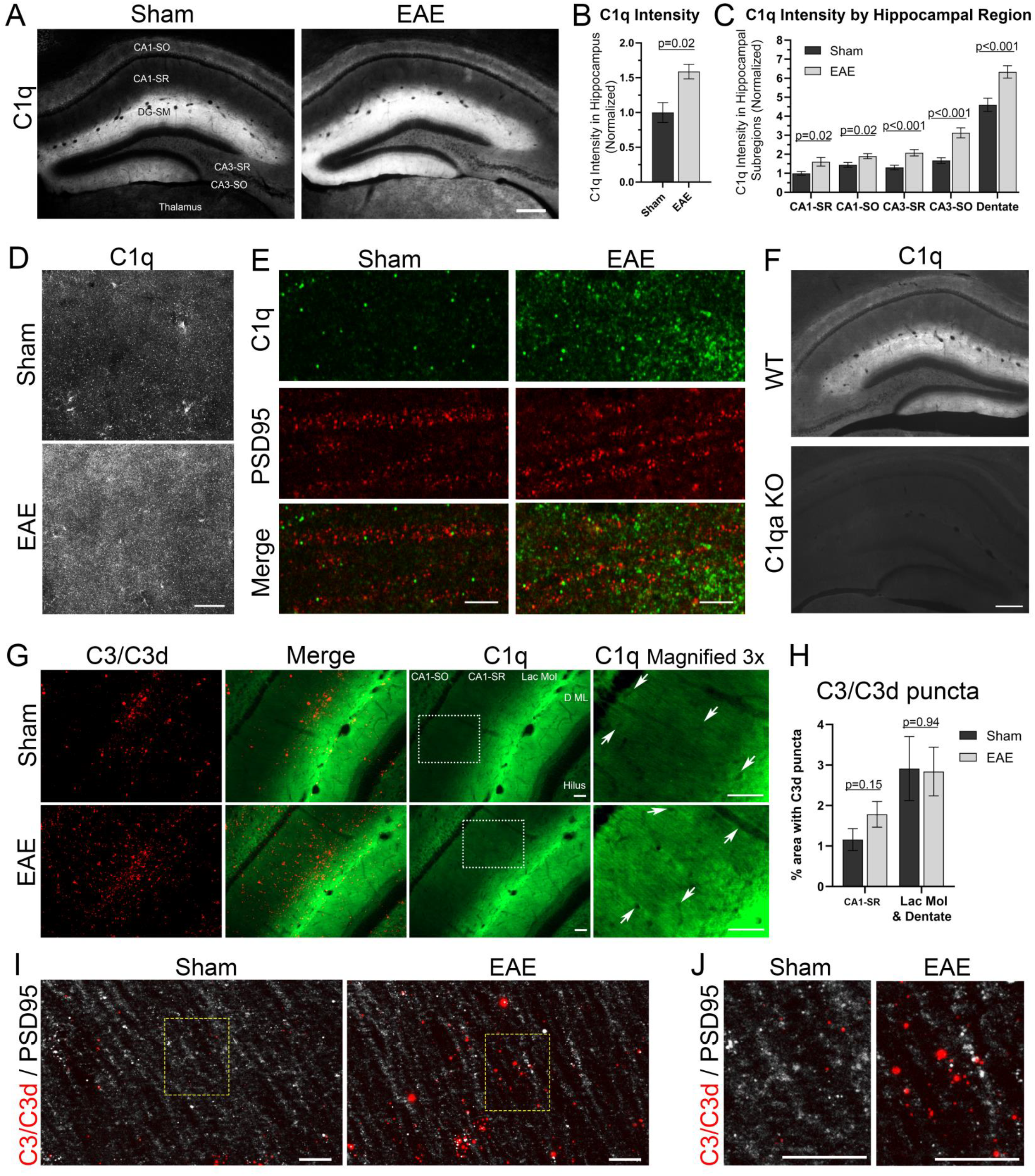
C1q and C3d protein levels are elevated in the hippocampus of EAE mice. A-D) Brain sections from EAE and Sham control mice were immuno-labeled for C1q. A) Images show C1q in the hippocampus; Scale bar = 250μm. B) Quantification of mean C1q fluorescence intensity in hippocampus normalized to sham controls. N=4 Statistical Analysis: Student’s t-test. C) Quantification of C1q intensity within hippocampal sub-regions. SR=Striatum Radiatum; SO=Striatum Oriens; DG=Dentate Gyrus. N=17 animals per group. Statistical Analysis: Student’s t-test for each region. D) Higher-magnification images show considerable diffuse as well as punctate C1q expression in CA1-Stratum Radiatum from Sham and EAE mice. Scale bar = 20μm. E) C1q and PSD95 immuno-stain in Sham and EAE brain sections within the CA1-Stratum Radiatum. Scale bar = 5μm. F) C1q antibody (Abcam Ab182451) is specific for C1q as no staining is visible in C1qa KO mice. Scale bar = 250 μm. G) C3/C3d and C1q immuno-staining in hippocampus of Sham and EAE. The C3d antibody recognizes full length C3 as well as any C3 cleaved fragments that contain the C3d region. Images on far right are 3x magnifications shown at increased brightness/contrast of boxed regions in the C1q images. Arrows highlight regions of higher C1q surrounding blood vessels. Lac Mol = Lacunosum Molecular; D ML=Dentate Molecular Layer. Scale bars = 60μm H) Quantification of C3d intensity within hippocampal sub-regions CA1-Striatum Radiatum and the C1q high expression region encompassing the Lacunosum molecular layer and Dentate Molecular layer. N=10 animals per group. Statistical Analysis: Student’s t-test for each region. I-J) C3/C3d and PSD95 in CA1-Stratum Radiatum at 60x magnification. Scale bars = 11 μm. Yellow boxes in I are enlarged in J. For all graphs in figure, error bars=SEM.

The C3d antibody, which detects both full-length C3 and C3 cleavage products that contain the C3d region reflects both expression and chronic deposition of the active complement pathway. C3/C3d puncta are found at a low level throughout the hippocampus with the highest density occurring in sham mice within the lacunosum molecular layer (Figure 2 F-G). In EAE mice, there was an increase in C3/C3d puncta in the CA1-stratum radiatum region of the hippocampus compared to sham that is just shy of significant (Figure 2 G; % of CA1-SR area occupied by C3/C3d: EAE: 1.8% vs Sham: 1.2%; p=0.15 *t*-test). The lacunosum and dentate molecular layers that have higher C3/C3d expression at baseline did not have increased C3/C3d puncta in EAE compared to sham. Like C1q, C3/C3d puncta in the CA1-SR occasionally co-localized to PSD95+ synapses and along dendrites, but the C3/C3d puncta are also found independent of PSD95 puncta (Figure 2 I-J).

### Comparison of EAE pathology in WT, C1qa KO, and C3 KO Mice Loss of C3 but not C1q results in less severe EAE motor impairment

Next, we immunized wild type (WT), C1qa knockout (KO) and C3 KO mice for EAE to determine whether the early complement pathway is a key driving component of synapse loss within the grey matter in EAE, particularly in the hippocampus. We also assessed whether the loss of C1q or C3 could protect against the EAE-induced motor impairment caused by spinal cord damage (Figure 3). As previously reported [33, 35, 38], C3 knockout resulted in less severe EAE motor deficits with a significant decrease in the mean clinical score both at peak disease (14-15 days post immunization) and during the chronic phase (20-30 days post immunization). On the other hand, C1qa KO did not alter the EAE disease course in any way as the C1qa KO EAE mean clinical scores were virtually identical to WT EAE. Neither C1qa KO nor C3 KO altered the timing of motor symptom onset. This result is consistent with the conclusion that it is primarily the alternative complement pathway that plays an important role in EAE white matter disease [29].

**Figure 3:**
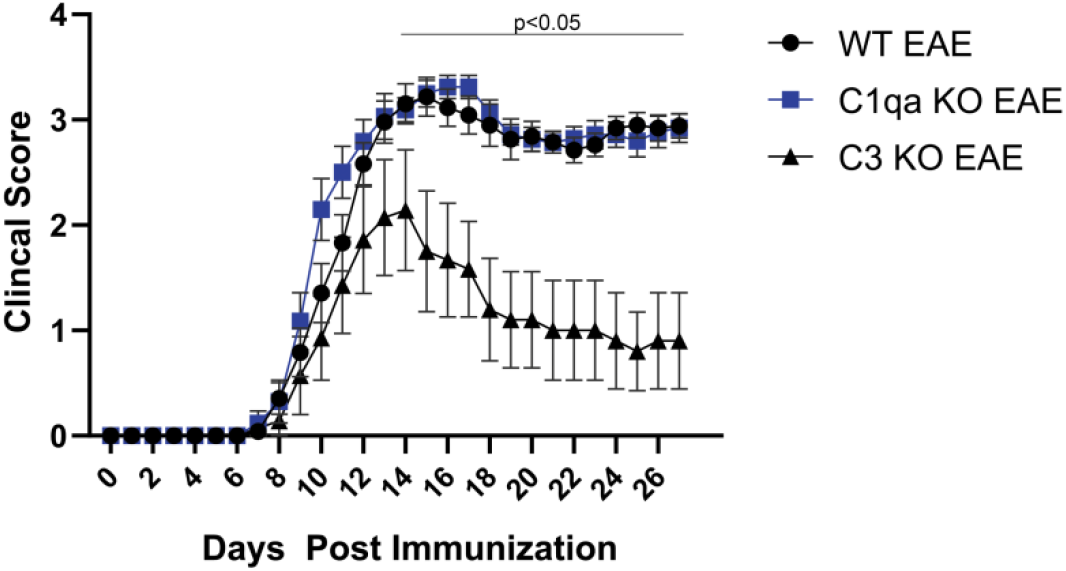
Loss of C3, but not C1q, resulted in a reduction in the mean severity of EAE motor deficits. Graph of the mean clinical scores from EAE immunized WT, C1qa KO, and C3 KO mice. WT: n=24; C1qa KO n=17; C3 KO n=7. Error bars=SEM. Statistical Analysis: Kruskal-Wallis test with Dunn’s multiple comparisons test. C3 KO EAE vs WT EAE has P<0.05 beginning on Day 14.

### C1qa and C3 knockout mice have less EAE-induced synapse loss

As we have previously documented significant synapse elimination in the CA1-stratum radiatum layer of the hippocampus of EAE mice, we chose to focus on this same region for this study assessing the role of complement in grey matter synapse loss. Loss of early complement genes C3 and C1qa protected synapses during EAE (Figure 4). WT mice with EAE displayed a 13% decrease in the number of Homer1 puncta in the CA1-striatum radiatum of the hippocampus compared to sham controls (p=0.003; Sidak). C1qa KO mice showed partial protection against EAE-induced synapse elimination as they had only 7% loss in Homer1 synaptic density compared to C1qa KO sham controls, and the synapse loss was no longer statistically significant (p=0.42, Sidak). C3 KO also were protected against EAE-induced synapse loss with no significant difference between sham and EAE (3% decrease; p=0.97). We found no differences in absolute synapse counts between WT, C1qa KO, and C3 KO mice that all had ~4000 synapses per 10μm image stack (WT: 3933 +/− 113; C1qa KO: 3951 +/− 118; C3 KO: 3929 +/− 175) showing that the genetic knockout of complement genes did not change the development of the hippocampus in a way that affected synapse numbers. Just as we saw that genetic loss of C3 and C1qa prevented loss of Homer1+ synapses due to EAE, we found PSD95+ synaptic puncta were likewise protected by loss of complement genes. WT EAE mice had a significant, 17% loss of PSD95+ postsynaptic puncta compared to sham controls (p=0.02, Sidak) whereas C1qa KO EAE mice had 11% fewer and C3 KO EAE mice had 6% fewer PSD95+ synaptic puncta than sham controls of the same genotype. Both C1qKO and C3 KO EAE groups were no longer significantly different from sham controls (C1qKO p=0.39; C3 KO p=0.94, Sidak). In summary, C1qa KOs with EAE showed a reduced loss in synaptic density over WT, suggesting that C1q plays a small role in EAE-induced synapse elimination but is not the primary driver. Knockout of C3 proved to be much more effective at preventing synapse loss. This could suggest that the alternative pathway is more important in grey matter pathology through C3-mediated opsonization and phagocytic removal and/or pro-inflammatory signaling via C3a.

**Figure 4:**
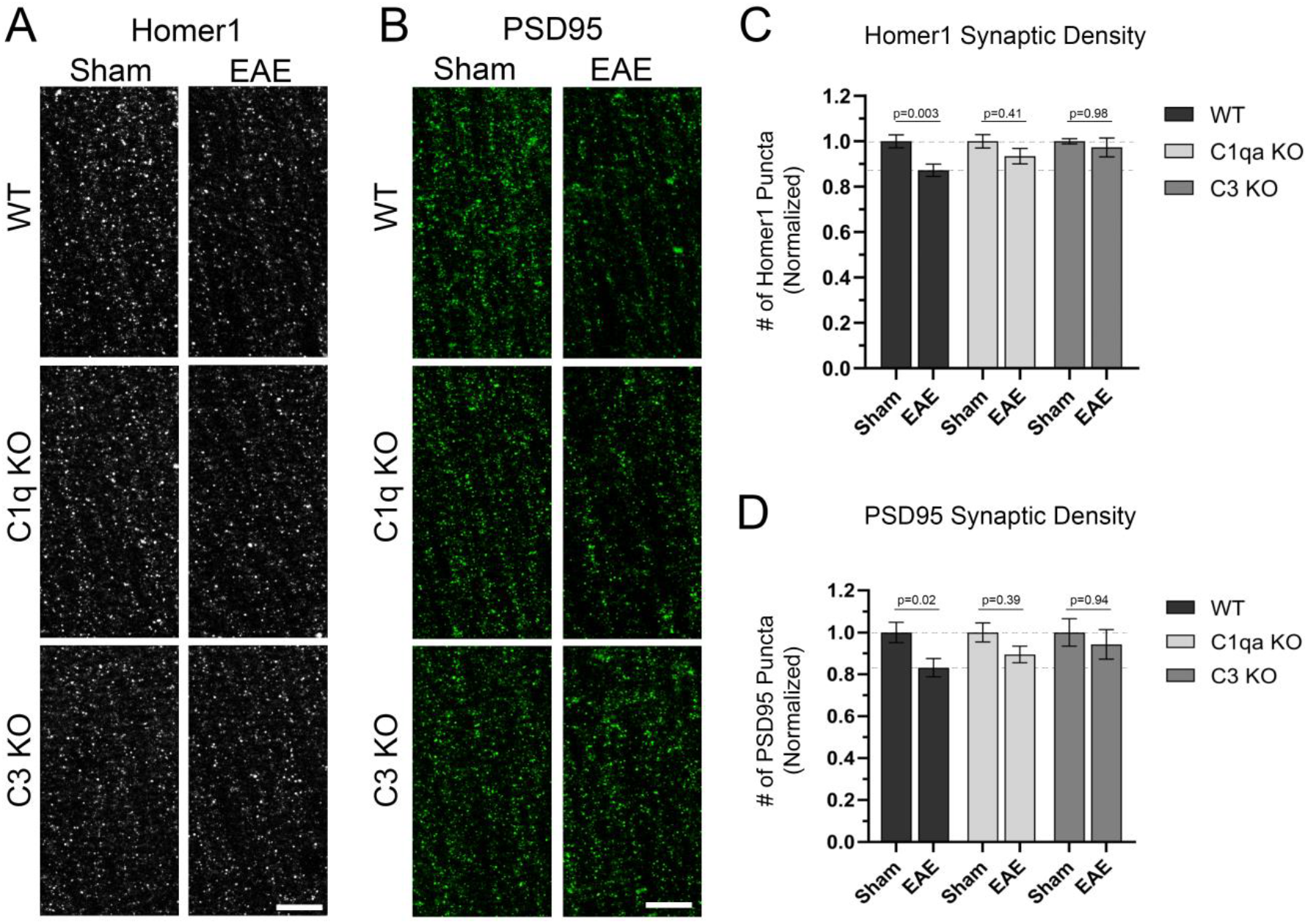
C1qa and C3 knockout mice have less EAE induced synapse loss in the CA1-SR region of the hippocampus. A-B) Brain sections from WT, C1qa KO, and C3 KO sham and EAE immunized mice were immuno-stained for the postsynaptic marker Homer1 (A) and PSD95 (B). Scale bars = 10μm. C) Quantification of Homer1 synaptic puncta density normalized to sham controls of each genotype. D) Quantification of PSD95 synaptic puncta density normalized to sham controls of each genotype. For data set: WT sham: n=23-25; WT EAE: n=19; C1qa KO sham: n= 13-14; C1qa KO EAE: n=14; C3 KO sham: n=6; C3 KO EAE: n=5. Statistical Analysis: ANOVAs with Sidak’s multiple comparison tests. Error bars=SEM.

### C3 knockout mice, but not C1q knockout mice, have less microglial activation

We next assessed whether C1qa or C3 KO could alter the morphometric parameters of microglial activation induced by EAE in the hippocampus. In WT mice, EAE immunization results in increased IBA1 expression by microglia as well as a change in microglial morphology that results in thicker and shorter processes that has been associated with a reactive microglial phenotype (Figure 5). This change in microglial cell morphology can be measured by a decrease in the surface area/volume ratio and a decrease in the skeletal length/volume ratio of IBA1+ cells. WT EAE mice have a significant 1.24-fold increase in the average IBA1+ cell volume per field of view compared to WT Sham (p=0.03; Sidak) and a significant 1.46-fold increase in the summated IBA1 intensity compared to WT Sham (p=0.003). C1qa KO EAE mice show a similar 1.38-fold increase in IBA1+ cell volume and a 1.57-fold increase in IBA1 intensity compared to C1qa KO shams (p=0.003 and p=0.004 respectively). However, C3 KO EAE mice had no increase in IBA1+ cell volume (1.01-fold; p=0.99) and only a small, insignificant increase in IBA intensity (1.14-fold; p=0.94) compared to C3 KO shams. Similar results were obtained for the surface value/volume ratio and skeletal length/volume ratio. C1qa KO EAE and WT EAE mice had similar significant decreases in these two microglial morphology measurements compared to genotype-matched sham controls (Surface area/volume: WT EAE 0.88, p<0.001; C1qa KO EAE 0.88, p=0.001 compared to 1.0 in genotyped matched sham controls; Skeletal Length/volume: WT EAE 0.77, p<0.001; C1qaKO EAE 0.76, p=0.002; compared to 1.0 in genotyped matched sham controls). C3 KO EAE mice, however, only showed small, insignificant decreases in the surface area/volume ratio (C3 EAE 0.96 vs C3 sham 1.0; p=0.68) and skeletal length/volume ratio (C3 KO EAE: 0.93 vs C3 sham 1.0; p=0.89). Taken together, these results show that C3 KO prevented hippocampal microglial activation induced by EAE, but C1qa KO had no effect on microglial morphology in EAE.

**Figure 5:**
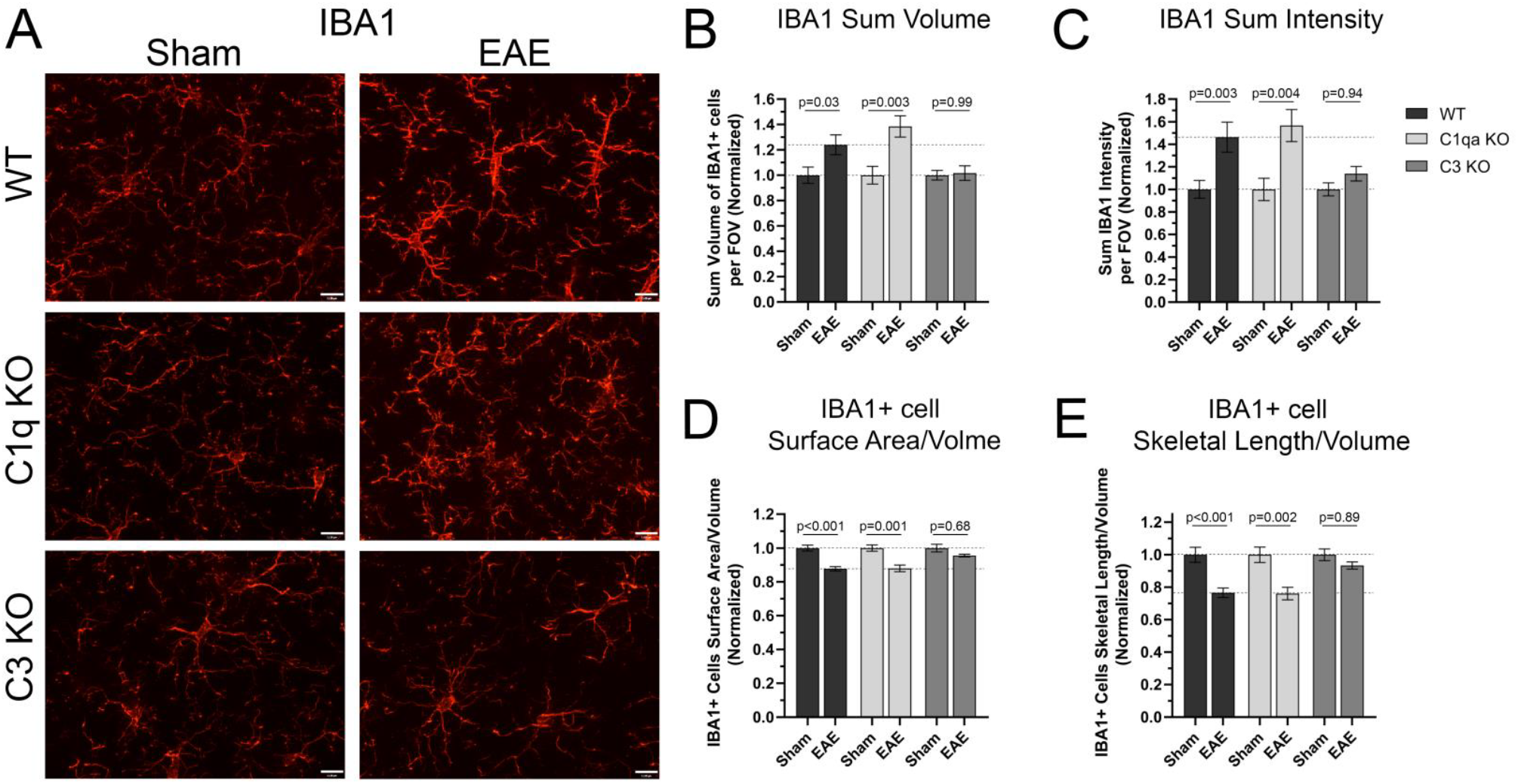
C3 knockout mice with EAE have reduced microglia activation compared to WT EAE mice. Loss of C1qa had little to no effect on microglia activation induced by EAE. A) Brain sections from WT, C1qa KO, and C3 KO sham and EAE immunized mice were immuno-stained for the microglia protein: IBA1 and imaged in the CA1-stratum radiatum of the hippocampus. Scale bars = 11μm. B-E) Quantification of microglia (IBA1+ cell) expression and morphology normalized to sham controls of each genotype. B) Quantification of the Sum Volume of IBA1+ cells per 60x field of view. C) Quantification of the Sum IBA1 Intensity per 60x field of view. D) Quantification of the IBA1+ cell surface area to volume ratio. E) Quantification of the IBA1+ cell skeletal length to volume ratio. For data set: WT sham: n=25; WT EAE: n=19; C1qa KO sham: n= 14; C1qa KO EAE: n=14; C3 KO sham: n=6; C3 KO EAE: n=5. Statistical Analysis: ANOVAs with Sidak’s multiple comparison tests. Error bars=SEM.

## Discussion

The overall goal of this study was to investigate complement expression and the contributions of complement proteins to synaptic damage and microglial activation in an EAE hippocampal model of MS grey matter pathology. Our results show that mice immunized with MOG_35-55_ to produce EAE have increased local expression of complement components C1q and C3 within the hippocampus. Specifically, C1q was elevated in all regions of the hippocampus and deposited throughout the neuropil, including some co-localization with synapses and dendrites. Increases in C3/C3d deposition were found in the CA1-stratum radiatum region of the hippocampus from EAE mice compared to sham controls. Knockout of C1q resulted in a small level of protection against EAE-induced synapse loss in the CA1-stratum radiatum region of the hippocampus. However, C1q knockout did not alter morphological markers of microglial activation, nor did it change EAE disease course measured by the clinical score. Knockout of C3 significantly attenuated the severity of EAE motor impairments and protected against synapse loss and microglial activation in the hippocampus. Thus, pathologic expression and activation of the early complement pathway drives a portion of the synapse elimination and microglial activation observed in EAE grey matter with C3 being a more important driver than C1q. This difference is not surprising as C3 deletion prevents most complement-mediated functions, whereas C1q deletion only blocks initiation by the classical pathway. It also suggests that the alternative pathway, in addition to the C1q-initiated classical pathway, contributes to synapse elimination and other EAE pathology.

Previous studies have shown that genetic ablation of alternative complement pathway components (C3, Factor B) or receptors to C3 cleavage products (C3aR, CR3 (CD11b-/-), CR4 (CD11c-/-) protects against EAE spinal cord pathology and associated motor deficits [33, 35-38, 55]. Additionally, loss of endogenous inhibitors like murine membrane-bound Crry exacerbate the EAE clinical score [56] and treatment with complement inhibitors that limit the C3 convertase, such as CR2-Crry, CR2-CFH, and sCR1, or antibodies that block the alternative pathway have been shown to attenuate the EAE clinical score [29, 57–59]. On the other hand, components of the classical pathway (C1q, C4) seem largely dispensable for MOG_35-55_-induced EAE motor impairments with genetic deletions resulting in little to no change in the EAE clinical score, although they may contribute to an auto-antibody exacerbation model of EAE [60, 61]. Our data corroborate this, as C1qa KO had no effect on the EAE clinical score whereas C3 KO mice were partially protected from EAE motor impairments.

This is the first study to investigate complement’s contribution to synaptic damage within EAE grey matter, where genetic deletion of C1q showed a small level of protection against EAE-induced synapse loss while knockout of C3 provided greater protection. Complement opsonization of synapses followed by microglia phagocytosis is a mechanism for synapse pruning during development and in some disease models including Alzheimer’s, frontotemporal dementia, viral encephalitis, aging, traumatic brain injury, and schizophrenia [11, 15-22, 62, 63]. Our results suggest that EAE (and perhaps multiple sclerosis) can be added to this list, although the precise mechanistic roles of C1q and especially C3 need additional studies to elucidate. As C3-dependent aspects of the complement pathway contributed not only to decreases in hippocampal synaptic density and microglia activation within the EAE hippocampus but also to motor deficits connected to white matter spinal cord pathology, it is possible that C3 KO may protect grey matter secondarily via systemic reduction in EAE severity in addition to direct effects within the grey matter. Thus it is difficult in this model system to distinguish between hippocampal protection due to loss of C3 activity directly at the synaptic level versus more global effects that result in wider reduction in disease severity. Yet, the fact that C1qa KO did not reduce the EAE clinical score, but moderately protected synapses suggests that some complement mediated synaptic pruning is locally occurring in the EAE hippocampus. Due to the many roles of complement in mediating immune cell functions, the role of C3 in EAE is likely complex. C3 is known to contribute to co-stimulation strength of T-cells and B-cells, chemo-attraction of infiltrating cells, cytokine production, opsonization of myelin and other debris, and likely opsonization of synapses [33, 35, 38, 64-66]. Unfortunately, there is no currently available conditional C3 KO mouse to probe the local role of C3 within the hippocampus and it will be important in the future to tease apart the role C3 plays in complement-dependent synaptic pruning from those in white matter inflammation and systemic immune activation in EAE. Regardless though, in light of evidence of complement activation and synaptic phagocytosis reported in grey matter from MS patients [9, 39], the protective effects of C3 KO and C1qa KO on EAE grey matter pathology are exciting given the paucity of treatments for grey matter neuroprotection in MS.

In summary, genetic loss of C1q or C3 provides protection from EAE induced synapse loss within grey matter while C3 deletion additionally protects against microglial activation. This coupled with the protection from EAE motor impairments, in the case of C3 KOs, suggest that the early complement pathway may be an important therapeutic target for MS and may be especially relevant to the grey matter neurodegeneration in progressive MS.

## Author Contributions

JWH, MJB, and HAG contributed to the study design. JWH and MJB performed all animal work. JWH, MJB, CW, WQQ, HL, SL, and YL did the tissue preparation, IHC staining, and image collection. JWH performed the western blots. JWH and PS did the qPCR experiments. JWH, MJB, and HAG contributed to data analysis and interpretation. JWH wrote the manuscript. All authors discussed, edited, and approved the manuscript.

## Acknowledgements

This work was funded by the National Multiple Sclerosis Society: NMSS RG-1607-25423 (MJB), the National Institutes of Health: 1R44NS092137 (HAG) and 1R21NS111255 (JWH); and the Harry T. Mangurian Jr. Foundation (JWH). The authors thank Dr. Andrea Tenner (University of California, Irvine) for providing the C1qa KO mice and the C1q antibody (1151). We also thank Angela Stout and Jeffrey M. Chamberlain for providing animal husbandry.

## Notes

Conflict of interest: The authors have declared that no conflict of interest exists.

## References

1. Coclitu, C., C.S. Constantinescu, and R. Tanasescu, The future of multiple sclerosis treatments. Expert Rev Neurother, 2016.

2. Calabrese, M., et al., Cortical lesion load associates with progression of disability in multiple sclerosis. Brain, 2012. 135(Pt 10): p. 2952–61.

3. Geurts, J.J., et al., Measurement and clinical effect of grey matter pathology in multiple sclerosis. Lancet Neurol, 2012. 11(12): p. 1082–92.

4. Rojas, J.I., et al., Brain atrophy in multiple sclerosis: therapeutic, cognitive and clinical impact. Arq Neuropsiquiatr, 2016. 74(3): p. 235–43.

5. Calabrese, M., et al., Exploring the origins of grey matter damage in multiple sclerosis. Nat Rev Neurosci, 2015. 16(3): p. 147–58.

6. Minagar, A., et al., The thalamus and multiple sclerosis: modern views on pathologic, imaging, and clinical aspects. Neurology, 2013. 80(2): p. 210–9.

7. Vercellino, M., et al., Grey matter pathology in multiple sclerosis. J Neuropathol Exp Neurol, 2005. 64(12): p. 1101–7.

8. Magliozzi, R., et al., A Gradient of neuronal loss and meningeal inflammation in multiple sclerosis. Ann Neurol, 2010. 68(4): p. 477–93.

9. Michailidou, I., et al., Complement C1q-C3-associated synaptic changes in multiple sclerosis hippocampus. Ann Neurol, 2015. 77(6): p. 1007–26.

10. Jurgens, T., et al., Reconstruction of single cortical projection neurons reveals primary spine loss in multiple sclerosis. Brain, 2016. 139(Pt 1): p. 39–46.

11. Stevens, B., et al., The classical complement cascade mediates CNS synapse elimination. Cell, 2007. 131(6): p. 1164–78.

12. Chung, W.S., et al., Astrocytes mediate synapse elimination through MEGF10 and MERTK pathways. Nature, 2013. 504(7480): p. 394–400.

13. Weinhard, L., et al., Microglia remodel synapses by presynaptic trogocytosis and spine head filopodia induction. Nat Commun, 2018. 9(1): p. 1228.

14. Paolicelli, R.C., et al., Synaptic pruning by microglia is necessary for normal brain development. Science, 2011. 333(6048): p. 1456–8.

15. Bialas, A.R. and B. Stevens, TGF-beta signaling regulates neuronal C1q expression and developmental synaptic refinement. Nat Neurosci, 2013. 16(12): p. 1773–82.

16. Vasek, M.J., et al., A complement-microglial axis drives synapse loss during virus-induced memory impairment. Nature, 2016. 534(7608): p. 538–43.

17. Lui, H., et al., Progranulin Deficiency Promotes Circuit-Specific Synaptic Pruning by Microglia via Complement Activation. Cell, 2016. 165(4): p. 921–35.

18. Shi, Q., et al., Complement C3-Deficient Mice Fail to Display Age-Related Hippocampal Decline. J Neurosci, 2015. 35(38): p. 13029–42.

19. Hong, S., et al., Complement and microglia mediate early synapse loss in Alzheimer mouse models. Science, 2016. 352(6286): p. 712–6.

20. Sekar, A., et al., Schizophrenia risk from complex variation of complement component 4. Nature, 2016. 530(7589): p. 177–83.

21. Fonseca, M.I., et al., Absence of C1q leads to less neuropathology in transgenic mouse models of Alzheimer’s disease. J Neurosci, 2004. 24(29): p. 6457–65.

22. Alawieh, A., et al., Identifying the role of complement in triggering neuroinflammation after traumatic brain injury. J Neurosci, 2018.

23. Hammond, J.W., et al., HIV Tat causes synapse loss in a mouse model of HIV-associated neurocognitive disorder that is independent of the classical complement cascade component C1q. Glia, 2018. 66(12): p. 2563–2574.

24. Lobsiger, C.S., et al., C1q induction and global complement pathway activation do not contribute to ALS toxicity in mutant SOD1 mice. Proc Natl Acad Sci U S A, 2013. 110(46): p. E4385–92.

25. Ingram, G., et al., Complement activation in multiple sclerosis plaques: an immunohistochemical analysis. Acta Neuropathol Commun, 2014. 2: p. 53.

26. Rus, H., C. Cudrici, and F. Niculescu, C5b-9 complement complex in autoimmune demyelination and multiple sclerosis: dual role in neuroinflammation and neuroprotection. Ann Med, 2005. 37(2): p. 97–104.

27. Weerth, S.H., et al., Complement C5 in experimental autoimmune encephalomyelitis (EAE) facilitates remyelination and prevents gliosis. Am J Pathol, 2003. 163(3): p. 1069–80.

28. Reiman, R., et al., Expression of C5a in the brain does not exacerbate experimental autoimmune encephalomyelitis. Neurosci Lett, 2005. 390(3): p. 134–8.

29. Barnum, S.R. and A.J. Szalai. Complement and demyelinating disease: no MAC needed? Brain Res Rev, 2006. 52(1): p. 58–68.

30. Reiman, R., et al., Disruption of the C5a receptor gene fails to protect against experimental allergic encephalomyelitis. Eur J Immunol, 2002. 32(4): p. 1157–63.

31. Mead, R.J., et al., The membrane attack complex of complement causes severe demyelination associated with acute axonal injury. J Immunol, 2002. 168(1): p. 458–65.

32. Michailidou, I., et al., Systemic inhibition of the membrane attack complex impedes neuroinflammation in chronic relapsing experimental autoimmune encephalomyelitis. Acta Neuropathol Commun, 2018. 6(1): p. 36.

33. Szalai, A.J., et al., Complement in experimental autoimmune encephalomyelitis revisited: C3 is required for development of maximal disease. Mol Immunol, 2007. 44(12): p. 3132–6.

34. Calida, D.M., et al., Cutting edge: C3, a key component of complement activation, is not required for the development of myelin oligodendrocyte glycoprotein peptide-induced experimental autoimmune encephalomyelitis in mice. J Immunol, 2001. 166(2): p. 723–6.

35. Smith, S.S., et al., Deletion of both ICAM-1 and C3 enhances severity of experimental autoimmune encephalomyelitis compared to C3-deficient mice. Neurosci Lett, 2008. 442(2): p. 158–60.

36. Bullard, D.C., et al., Critical requirement of CD11b (Mac-1) on T cells and accessory cells for development of experimental autoimmune encephalomyelitis. J Immunol, 2005. 175(10): p. 6327–33.

37. Bullard, D.C., et al., p150/95 (CD11c/CD18) expression is required for the development of experimental autoimmune encephalomyelitis. Am J Pathol, 2007. 170(6): p. 2001–8.

38. Nataf, S., et al., Attenuation of experimental autoimmune demyelination in complement-deficient mice. J Immunol, 2000. 165(10): p. 5867–73.

39. Watkins, L.M., et al., Complement is activated in progressive multiple sclerosis cortical grey matter lesions. J Neuroinflammation, 2016. 13(1): p. 161.

40. Bellizzi, M.J., et al., Platelet-Activating Factor Receptors Mediate Excitatory Postsynaptic Hippocampal Injury in Experimental Autoimmune Encephalomyelitis. J Neurosci, 2016. 36(4): p. 1336–46.

41. Ziehn, M.O., et al., Hippocampal CA1 atrophy and synaptic loss during experimental autoimmune encephalomyelitis, EAE. Lab Invest, 2010. 90(5): p. 774–86.

42. Ziehn, M.O., et al., Estriol preserves synaptic transmission in the hippocampus during autoimmune demyelinating disease. Lab Invest, 2012. 92(8): p. 1234–45.

43. Mandolesi, G., et al., Cognitive deficits in experimental autoimmune encephalomyelitis: neuroinflammation and synaptic degeneration. Neurol Sci, 2010. 31(Suppl 2): p. S255–9.

44. Nistico, R., et al., Inflammation subverts hippocampal synaptic plasticity in experimental multiple sclerosis. PLoS One, 2013. 8(1): p. e54666.

45. Kyran, E.L., et al., Multiple pathological mechanisms contribute to hippocampal damage in the experimental autoimmune encephalomyelitis model of multiple sclerosis. Neuroreport, 2018. 29(1): p. 19–24.

46. Hamilton, A.M., et al., Central nervous system targeted autoimmunity causes regional atrophy: a 9.4T MRI study of the EAE mouse model of Multiple Sclerosis. Sci Rep, 2019. 9(1): p. 8488.

47. Habbas, S., et al., Neuroinflammatory TNFalpha Impairs Memory via Astrocyte Signaling. Cell, 2015. 163(7): p. 1730–41.

48. Novkovic, T., et al., Hippocampal function is compromised in an animal model of multiple sclerosis. Neuroscience, 2015. 309: p. 100–12.

49. Planche, V., et al., Selective dentate gyrus disruption causes memory impairment at the early stage of experimental multiple sclerosis. Brain Behav Immun, 2017. 60: p. 240–254.

50. Wessels, M.R., et al., Studies of group B streptococcal infection in mice deficient in complement component C3 or C4 demonstrate an essential role for complement in both innate and acquired immunity. Proc Natl Acad Sci U S A, 1995. 92(25): p. 11490–4.

51. Botto, M., et al., Homozygous C1q deficiency causes glomerulonephritis associated with multiple apoptotic bodies. Nat Genet, 1998. 19(1): p. 56–9.

52. Huang, J., et al., Neuronal protection in stroke by an sLex-glycosylated complement inhibitory protein. Science, 1999. 285(5427): p. 595–9.

53. Bellizzi, M.J., et al., The Mixed-Lineage Kinase Inhibitor URMC-099 Protects Hippocampal Synapses in Experimental Autoimmune Encephalomyelitis. eNeuro, 2018. 5(6).

54. Fonseca, M.I., et al., Cell-specific deletion of C1qa identifies microglia as the dominant source of C1q in mouse brain. J Neuroinflammation, 2017. 14(1): p. 48.

55. Boos, L., et al., Deletion of the complement anaphylatoxin C3a receptor attenuates, whereas ectopic expression of C3a in the brain exacerbates, experimental autoimmune encephalomyelitis. J Immunol, 2004. 173(7): p. 4708–14.

56. Ramaglia, V., et al., C3-dependent mechanism of microglial priming relevant to multiple sclerosis. Proc Natl Acad Sci U S A, 2012. 109(3): p. 965–70.

57. Hu, X., S. Tomlinson, and S.R. Barnum, Targeted inhibition of complement using complement receptor 2-conjugated inhibitors attenuates EAE. Neurosci Lett, 2012. 531(1): p. 35–9.

58. Piddlesden, S.J., et al., Soluble recombinant complement receptor 1 inhibits inflammation and demyelination in antibody-mediated demyelinating experimental allergic encephalomyelitis. J Immunol, 1994. 152(11): p. 5477–84.

59. Hu, X., et al., Therapeutic inhibition of the alternative complement pathway attenuates chronic EAE. Mol Immunol, 2013. 54(3-4): p. 302–8.

60. Urich, E., et al., Autoantibody-mediated demyelination depends on complement activation but not activatory Fc-receptors. Proc Natl Acad Sci U S A, 2006. 103(49): p. 18697–702.

61. Boos, L.A., A.J. Szalai, and S.R. Barnum, Murine complement C4 is not required for experimental autoimmune encephalomyelitis. Glia, 2005. 49(1): p. 158–60.

62. Chu, Y., et al., Enhanced synaptic connectivity and epilepsy in C1q knockout mice. Proc Natl Acad Sci U S A, 2010. 107(17): p. 7975–80.

63. Schafer, D.P., et al., Microglia sculpt postnatal neural circuits in an activity and complement-dependent manner. Neuron, 2012. 74(4): p. 691–705.

64. Clarke, E.V. and A.J. Tenner. Complement modulation of T cell immune responses during homeostasis and disease. J Leukoc Biol, 2014. 96(5): p. 745–56.

65. Carroll, M.C. and D.E. Isenman, Regulation of humoral immunity by complement. Immunity, 2012. 37(2): p. 199–207.

66. van der Laan, L.J., et al., Macrophage phagocytosis of myelin in vitro determined by flow cytometry: phagocytosis is mediated by CR3 and induces production of tumor necrosis factor-alpha and nitric oxide. J Neuroimmunol, 1996. 70(2): p. 145–52.

